# Evidence of macroevolutionary selection of the mutation rate in sexual species

**DOI:** 10.1101/2023.07.16.549230

**Authors:** Gordon Irlam

**Affiliations:** Los Altos, California, United States

**Keywords:** population-optimal mutation rate

## Abstract

For sexual species it has been hypothesized that, over macroevolutionary timescales, species-level processes such as differential extinction may bias the germline mutation rates of surviving species toward values that maximize long-term population mean fitness. If this hypothesis is correct, then the mutation rate is expected to lie near a population-optimal value that depends on several evolutionary and demographic parameters. Using previously published data, these parameters were estimated for several well-studied species pairs, enabling a quantitative evaluation of this prediction. Across the species examined, empirical estimates of mutation rates fall within the range that is predicted by the underlying model to yield appreciable population mean fitness, given substantial uncertainty in several parameter values. The underlying model is consistent with a previously reported inverse relationship between effective population sizes and mutation rates within broad clades. Although not intended to provide precise predictions for individual species, these results are compatible with the hypothesis that macroevolutionary processes contribute to shaping germline mutation rates toward population-optimal values.

## Introduction

This manuscript empirically assesses the determinants of germline spontaneous mutation rates in sexual eukaryotes.

Because most mutations are harmful rather than beneficial, adaptations that reduce the mutation rate are expected to be favored by microevolution. Despite the expectation that microevolution acts to minimize the mutation rate, spontaneous mutations cause a significant genetic load; the mutational load [1]. Why, then, is the mutation rate not lower than observed?

The drift-barrier hypothesis argues that genetic drift acts to create a barrier that impedes the fixation of mutations that contribute only small improvements in replication fidelity [2, 3].

What is beneficial at the level of genes or organisms is not necessarily optimal at the population or species level. If the mutation rate is driven too low, the species may be unable to adapt sufficiently and will face an elevated risk of extinction. Too high, and the species will bear a large deleterious mutational load. There therefore exists a population-optimal mutation rate between these two extremes that maximizes long-term geometric mean population mean fitness. For sexual species it is possible to estimate this population-optimal mutation rate [4]. Since microevolution might largely favor smaller and smaller mutation rates, one plausible mechanism for species to achieve the population-optimal mutation rate is through macroevolution. Species with mutation rates that are too high or too low are more likely to go extinct, leaving surviving species biased towards mutation rates near the population-optimal value. This is not a new argument; it was articulated by Kimura in 1967 [5]. For brevity, this hypothesis might be termed the mutation rate macroevolutionary selection hypothesis.

This manuscript seeks to evaluate the mutation rate macroevolutionary selection hypothesis. For the three animal species pairs considered, the predicted population-optimal mutation rates are found to be close to the observed mutation rates. That is, both are around 10^*−*8^ mutations per base pair per generation. Apart from one of the species pairs, where the a priori parameters might have been misestimated, the observed mutation rate is found to lie within the small range of values that are expected to yield an appreciable population mean fitness. Assuming species tend towards the population-optimal mutation rate, a quantitative relationship between effective population size and the mutation rate is predicted. This relationship is shown to largely agree with a previously observed empirical relationship between these two variables. These observations add evidence to the hypothesis that the mutation rates of surviving species are biased by macroevolutionary processes towards values that produce optimal population mean fitness.

## Results

### The rate of adaptive substitution

Let Γ_*s*_ denote the per sexual generation rate at which substituting sites become established in the population. For a species pair over macroevolutionary timescales, the temporal mean value of Γ_*s*_ equals the rate of positively selected adaptive substitution per generation and can be estimated by[4],

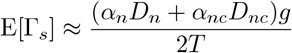

where *α*_*n*_ is the fraction of non-synonymous substitutions that are fixed by positive selection, *D*_*n*_ is the number of non-synonymous substitutions, *α*_*nc*_ is the fraction of non-coding substitutions that are fixed by positive selection, *D*_*nc*_ is the number of non-coding substitutions, *g* is the generation length, and *T* is the divergence time.

Table 1 shows the parameters used for, and estimates of, Γ_*s*_ for three species pairs. The *Drosophila* species studied were *D. melanogaster* and *D. simulans*. Although Γ_*s*_ is larger for humans and chimps than the two other species pairs, the much longer generation length results in a substantially lower per generation rate of genomic adaptation to environmental change. The value for *Drosophila* is broadly in line with an estimate of one coding adaptive substitution every 45 years for *D. simulans* and *D. yakuba* based on an analysis of just 35 genes [6].

**Table 1:** Estimation of Γ_*s*_, the rate at which substituting sites become established per sexual generation. Analysis of three species pairs. *Drosophila* is *D. melanogaster* and *D. simulans*

|  | human-chimp | mouse-rat | *Drosophila* | Comments |
| --- | --- | --- | --- | --- |
| $\alpha_n$ | 0.25 [7] | 0.41 [7] | 0.54 [8] | |
| $D_n$ | $3.9 \times 10^4$ [9] | $9.7 \times 10^5$ | $5.9 \times 10^4$ [8, 10] | mouse-rat: 3.5% [11]<br>$\times 75\%$ [12] $\times 3.7 \times 10^7$ [10] |
| $\alpha_{nc} D_{nc}$ | 0 [13] | $\geq 0$ | $1.3 \times 10^5$ [8, 10] | mouse-rat: not well known |
| $g$ | 26 [14, 15] | 0.3 [16, 17] | 0.07 [18, 19] | in years |
| $T$ | $6.4 \times 10^6$ | $1.2 \times 10^7$ | $4.6 \times 10^6$ | in years [20] |
| $E[\Gamma_s]$ | $2.0 \times 10^{-2}$ | $\geq 5.0 \times 10^{-3}$ | $1.2 \times 10^{-3}$ | |
| $\frac{E[\Gamma_s]}{g}$ | $7.7 \times 10^{-4}$ | $\geq 1.7 \times 10^{-2}$ | $1.8 \times 10^{-2}$ | |

### The population-optimal mutation rate

Let *µ* denote the germline spontaneous mutation rate per base pair per sexual generation. The population-optimal mutation rate 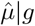 is expected to be given by [4],

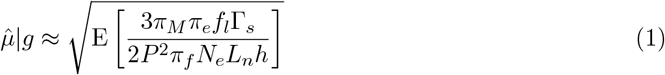

where E[*·*] denotes the expectation taken over generations, *π*_*M*_ is an adjustment to log fitness of the latent load (i.e., the genetic load from positively selected sites that are not yet established) due to macroevolutionary selection favoring fitter species, *π*_*e*_ is an adjustment to the log fitness effect of selection due to epistasis, *f*_*l*_ is the fraction of new positively selected sites that are latent (i.e., not yet established) sites, *P* is the ploidy (haploids 1, diploids 2), *π*_*f*_ is an adjustment to the probability of fixation at new positively selected sites as a result of unaccounted for interference between positively selected sites, *N*_*e*_ is the neutral site variance effective population size, *L*_*n*_ is the effective number of conserved sites in a haploid genome, and *h* is the harmonic mean value of the beneficial alleles’ dominance coefficients.

The given *g* qualifier in 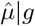 signifies that this is the optimal mutation rate for a given generation length. This may be different from the optimal mutation rate if *g* is also free to vary.

Over evolutionary time frames of the order of hundreds of millions of years, most terms in equation 1 are likely to be approximately constant. The exceptions are Γ_*s*_, *N*_*e*_, and *L*_*n*_. This means that for constant *N*_*e*_ and *L*_*n*_ or for constant Γ_*s*_, the expectation operator can be dropped and *N*_*e*_*L*_*n*_ represented by its harmonic mean value or Γ_*s*_ represented by its arithmetic mean value.

Equation 1 is intended for broad macroevolutionary inference and is unlikely to yield precise predictions for individual species. This is because other stochastic factors also affect the survival of species.

*N*_*e*_ was approximated by the current neutral site coalescent effective population size, 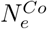. Unfortunately 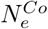 is only approximately appropriate over the time to the coalescent, not over the entire time until the species merged. Ideally *N*_*e*_ might be estimated using the pairwise sequentially Markovian coalescent model [21], and the generationally weighted harmonic mean value taken. Since this was not done, it is assumed that the variance effective population size over the entire time interval is approximated by the estimated coalescent effective population size. This approximation biases estimates toward recent demographic history.

Table 2 shows actual mutation rate *µ*, the parameters used for, and estimates of, 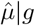 assuming a constant *N*_*e*_. In order to estimate the impact of macroevolutionary selection, *π*_*M*_, it was necessary to estimate *S*_*niche*_, the number of species occupying a given niche, *λ*, the rate of speciation along a lineage, and 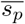, the mean selection coefficient of new positively selected sites. 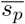 itself was determined from the mean fitness effects of new beneficial mutations, 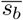.Macroevolutionary selection was then determined to have little impact; *π*_*M*_ ranged from 0.90 to 1.0. *L*_*n*_ was computed as the product of *f*_*n*_, the effective fraction of conserved sites in the genome, and *L*, the length of the genome. The final estimate of 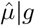 for mice and rats is a lower bound since the number of non-coding adaptive substitutions, *α*_*nc*_*D*_*nc*_, could not be reliably determined.

**Table 2:**
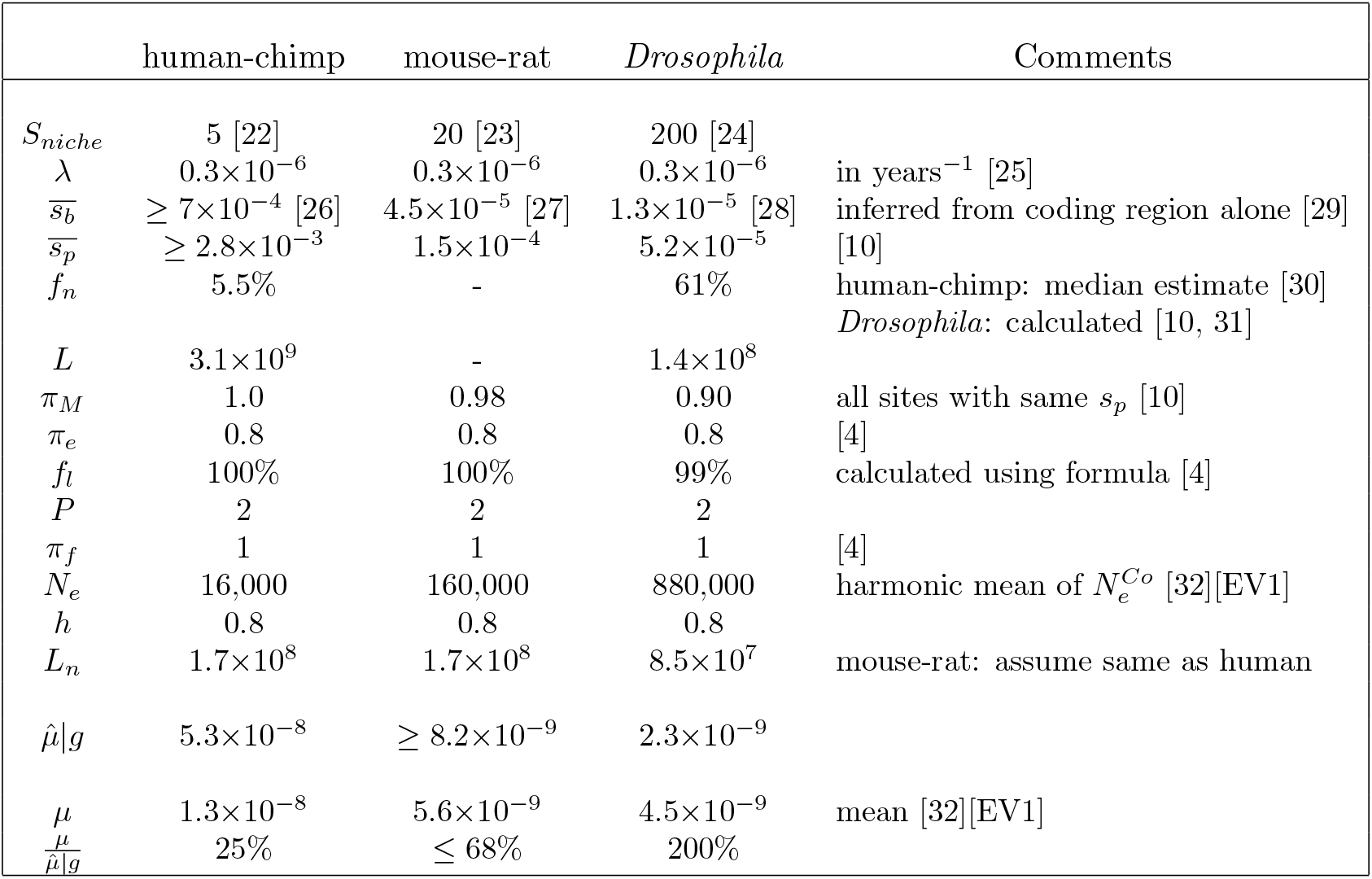
Observed mutation rates are reasonably close to hypothesized population-optimal mutation rates. Parameters values and a priori estimates of the population-optimal mutation rate, 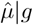, assuming a constant *N*_*e*_, along with the observed mutation rate *µ*, for three species pairs.

The values for the parameters in Table 2 were determined a priori, without reference to the resulting population-optimal mutation rate predictions. The predicted population-optimal mutation rates are reasonably close to the observed rates. I.e. both are around 10^*−*8^ mutations per base pair per sexual generation, and are at or within a factor of 4 of one another. Given uncertainties in several parameter values and the influence of additional stochastic factors, this correspondence provides supportive, though not definitive, evidence for the mutation rate macroevolutionary selection hypothesis.

### Post hoc considerations of the difference between the population-optimal and actual mutation rates

The estimated value of 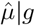 exceeds *µ* for humans and chimps, and for mice and rats, but is lower than *µ* for *Drosophila*.

Although exact concordance is not expected to occur, taken together, the factors listed below appear sufficient to allow 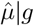 and *µ* to be in close agreement for each species pair. The uncertainty in the parameter values precludes determining whether *µ* systematically lags or exceeds 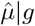.

### Humans and chimps

For humans and chimps, the estimate *α*_*n*_ = 0.25 appears high. Other estimates of *α*_*n*_ include 0.24 [33], 0.1–0.2 [34], 0.14 [35], 0.10–0.13 [36], 0.002 [37], 0.0 for human-macaque with a 95% confidence interval [-0.30, 0.24] [38], and *<* 0 [39]. Reducing *α*_*n*_ would reduce 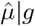. For example, using *α*_*n*_ = 0.1 would reduce 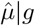 by 37%.

The ancestral human-chimp effective population size is estimated to have been 48,000 [40]. This is considerably larger than the estimate of 16,000 based on short-term coalescence that was used. A larger *N*_*e*_ would reduce 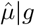. For example, using the ancestral estimate for *N*_*e*_ would reduce 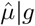 by 42%.

### Mice and rats

Mice and rats have a larger *N*_*e*_ than humans. This makes alleles that would be nearly neutral in humans, selected in mice and rats. This applies to regulatory elements in the non-coding region [41]. Mice and rats will thus have a greater number of effectively conserved sites than humans, *L*_*n*_. *L*_*n*_ for mice and rats was assumed to be the same as for humans. Increasing it will reduce 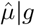. For example, doubling *L*_*n*_ would reduce 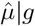 for mice and rats by 29%.

### Drosophila

The mutation rate in Drosophila was estimated as 4.5*×*10^*−*9^. This is the mean of 6 studies ranging from 1.7*×*10^*−*9^ to 6.6*×*10^*−*9^ [32][EV1]. If the historical actual value is towards the low end of the estimates from these studies, then *µ* and 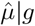 would be in agreement for Drosophila.

The time of divergence used for *Drosophila* was 4.6*×*10^6^ years ago. The 95% confidence interval is [2.5*×*10^6^, 1.3*×*10^7^] years ago [20], a considerable range. If the actual time was towards the lower end of this range, this would increase 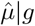. For example, using the lower interval value would increase 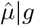 by 36%.

The genus *Drosophila* contains around 2,000 species that are known to have diversified rapidly [42]. Hawaiian *Drosophila* are a clade of an estimated 1,000 species that arose through founder events and vicariance, and form an adaptive radiation that began 26 million years ago [43]. This suggests that over much of the timespan of the genus *Drosophila, N*_*e*_ might have been influenced by founder events, and so be much lower than the estimate of 880,000 for *D. melanogaster* and *D. simulans*. Consequently, depending on the time frame over which macroevolutionary selection influences the mutation rate of surviving species, the observed mutation rate for *D. melanogaster* and *D. simulans* may be more appropriate to species with a lower *N*_*e*_. Reducing *N*_*e*_ would increase 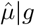.

### Predicted fitness

Let 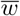 denote the population mean relative fitness of a sexual population. The geometric-mean value of 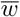 averaged over the number of generations as a function of *µ* is proposed to be given by [4],

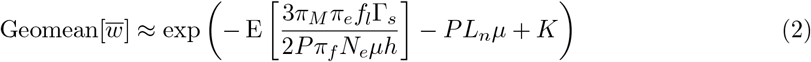

where *K* is either independent of, or only weakly dependent on, *µ*.

Using equation 2, Figure 1 illustrates Geomean[*w*] as a function of *µ* assuming constant *N*_*e*_, along with the estimated actual mutation rate. Figure 1 shows that there is a reasonably narrow range of mutation rates, within a factor three, outside of which population mean fitness is substantially smaller than at the population-optimal mutation rate. Equation 2 for geometric-mean population mean fitness implies a mutation rate “filter” through which competing species must pass. Except for humans and chimps, where *α*_*n*_ and *N*_*e*_ might have been misestimated, the mutation rates occurred where the species were expected to experience appreciable fitness. For humans and chimps a post hoc analysis with *α*_*n*_ = 0.1 and *N*_*e*_ = 48,000 showed the mutation rate occurred where it was expected to yield appreciable fitness. Such concordances would be more difficult to explain if the mutation rate macroevolutionary selection hypothesis were incorrect.

**Figure 1:**
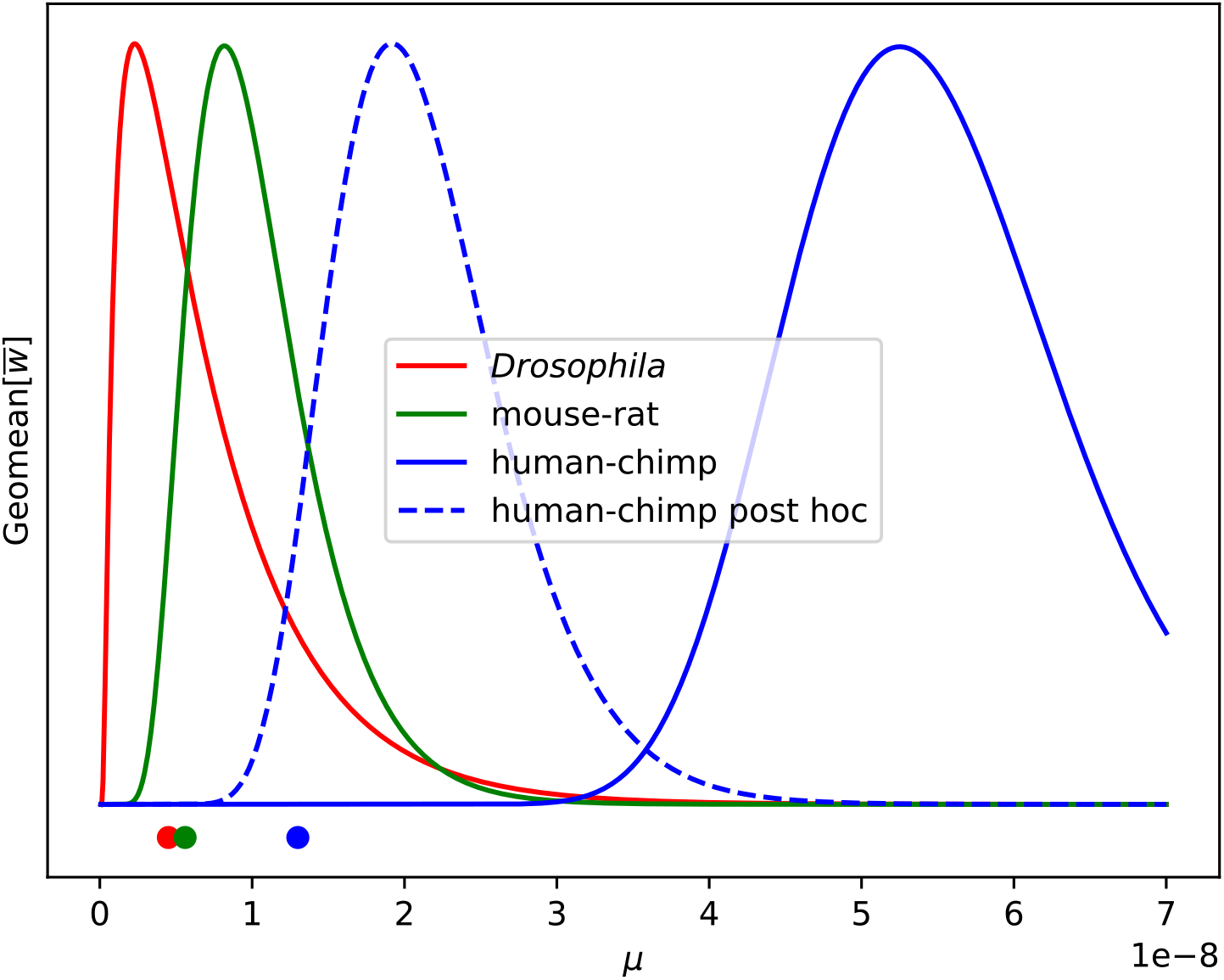
Population mean fitness implies reasonably narrow ranges for mutation rates. Generationally-averaged population mean fitness Geomean[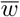] assuming a constant *N*_*e*_, for a priori parameter values as a function of the mutation rate *µ* (solid lines), for post hoc parameter values (dashed line), and observed mutation rates (dots). The mouse-rat curve is the a priori lower bound.

### The relationship between effective population size and the mutation rate

*π*_*M*_, *π*_*e*_, *π*_*f*_, *f*_*l*_, *P*, and *h* are expected to have similar values within very broad clades (Linnaean kingdoms). Much of the time *L*_*n*_ might also be expected to have roughly similar values within slightly less broad clades (Linnaean phyla). If either *N*_*e*_ or Γ_*s*_ is constant over time for individual species, then equation 1 implies that within a less broad clade,

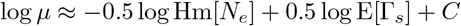

where Hm[*·*] denotes the harmonic mean taken over generations, and *C* is largely constant for the group of species.

Empirically, Lynch et al. report an approximately linear negative relationship between the log coalescent effective population size and the log mutation rate within broad clades of species (Linnaean phyla) [32]. The slopes of the log-log relationships ranged from −0.32 to −1.53 [32][Appendix S1].

Variability in Γ_*s*_ and possibly *L*_*n*_ between species presumably adds noise to the relationship. Lynch et al.’s finding of an inverse log-log relationship between *N*_*e*_ and *µ* counts as potential evidence for the mutation rate macroevolutionary selection hypothesis. It should be noted that the drift-barrier hypothesis also predicts an inverse relationship between *N*_*e*_ and *µ*, but does not uniquely predict the functional form or slope of the relationship [3].

## Discussion

### How might the mutation rate per sexual generation increase?

Proposing convergence toward a population-optimal mutation rate requires an accompanying biological mechanism for adjusting that rate. When the mutation rate is higher than optimal, microevolutionary forces can be invoked to bring it down. When the rate is lower than optimal it might appear that the only possibility is for a macroevolutionary force favoring extinction to take hold. However, this need not be the case.

One possibility is for a new species to simultaneously acquire a higher mutation rate and higher population-mean fitness through stochastic species founder effects. The scope for such a mechanism appears limited.

A more likely option appears to be through an increase in the generation length. Suppose substituting sites arise in response to changes in the external environment. Then Γ_*s*_ is given by the product of a rate per unit time, 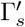, which is determined by the external environment, and the generation length, *g*. From equation 1, the population-optimal mutation rate per sexual generation is then,

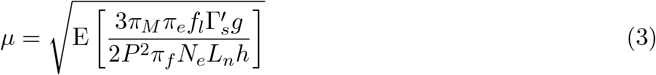

For most species, microevolution favors an increase in the generation length; the longer the generation length of an organism, the greater the number of offspring. By equation 3, increasing the generation length will increase the population-optimal mutation rate. At first glance, this seems to be the opposite of what is required. The mutation rate is too low, and the population-optimal mutation rate is made larger. However, at least in some species, when the generation length increases, there is a biological mechanism that increases the germline spontaneous mutation rate. This mechanism is the stem cell cycle that leads to the production of new germ cells [44]. Thus the mutation rate increases in a more or less linear fashion as the generation length increases, whereas the population-optimal mutation rate increases at the square root of the generation length. Consequently, microevolutionary forces favoring increased generation lengths can elevate the mutation rate, and if initially low, potentially bring it closer to its population-optimal value.

The stem cell cycle occurs in men and male mice, and in both sexes in *Drosophila melanogaster* [44]. With each stem cell division there is the chance for mutations to occur as the DNA is replicated.

It is hypothesized that something akin to the stem cell cycle in which mutation rates increase in proportion to the sexual generation time is widespread among sexual eukaryotes. Otherwise it would appear that the only way for mutation rates to increase would be through stochastic species founder effects.

## Conclusion

In this manuscript, the possibility that evolution drives the spontaneous mutation rate towards its population-optimal value was tested. When parameterized using independently estimated quantities, the mutation rate macroevolutionary selection hypothesis predicts the order of magnitude value of *µ*, an inverse relationship between *µ* and *N*_*e*_, the log–log form of this relationship, and its approximate slope. Together, these results are compatible with the hypothesis that differential macroevolutionary selection biases the spontaneous mutation rate of surviving species toward the value that maximizes long-term population mean fitness.

## Materials and methods

Assuming it exists, the variance effective population size, *N*_*e*_, can be approximated by the eigenvalue effective population size, 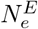 [45]. 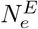 approaches the coalescent effective population size, 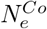 as population size increases provided that 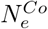 exists [46]. Thus *N*_*e*_ can be estimated using 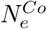, which is determined from the heterozygosity of neutral sites in the genome.

In order to determine *π*_*M*_ a simulation of the essential elements of macroevolutionary selection over time was performed. Fitness of a species was determined by the interplay between the stochastic arrival of latent sites, and their becoming substituting sites with fixed probability per unit time. At stochastic intervals a random species was cloned and replaced the then least fit species. All latent sites were assumed to share the same selection coefficient, equal to the mean value of the expected distribution of fitness effects for latent sites. This highly simplified model was intended only to estimate *π*_*M*_.

The number of species occupying a niche, *S*_*niche*_, was estimated as the average number of observed overlapping similar species (i.e., anthropoids, rodents, or drosophilids) in the wild. This value reflects both a reduction, since not all of the overlapping species are likely in competition, and an increase, on account of competition from more distantly related species.

The distribution of fitness effects for new positively selected sites was assumed to be exponential with mean 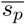 [47]. It differs from the distribution of fitness effects of beneficial mutations because beneficial mutations of small fitness effect are less likely to establish and fix than beneficial mutations of larger effect, and thus they will reoccur more frequently. The distribution of fitness effects of beneficial mutations is frequently studied. A mapping from the fitness effects of beneficial mutations to 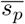 was thus sought.

Let *ρ*_*b*_(*s*) denote the distribution of fitness effects for new beneficial mutations for selection coefficient *s*, and 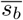 be the mean selection coefficient of effective beneficial mutations. Consider a particular value for the selection coefficient of new positively selected sites, *s*. Since *s* is exponentially distributed with mean 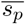, and the probability of fixation is approximately 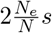 where *N* is the census population size [48, 49],

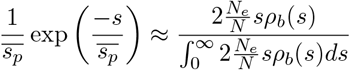

which implies that,

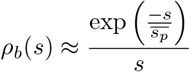

For positively selected sites with beneficial alleles having selection coefficients less than 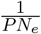 drift dominates over selection, and a beneficial mutation at the site is unlikely to contribute appreciably to the latent load, nor to the mutational load. Thus the mean selection coefficient for effective beneficial mutations is approximately given by,

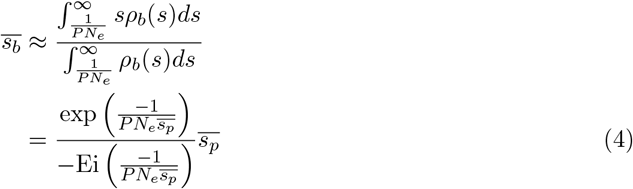

where Ei is the exponential integral,

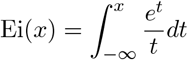

Equation 4 can then be solved for 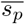 using standard numerical root finding techniques.

*h* for deleterious alleles is typically around 0.2 or 0.3 [50], suggesting a value of *h* for beneficial alleles equal to 1 minus this value, or around 0.8.

## Data availability

The code and data used in this study are publicly available as File S1. File S1 - Population-optimal mutation rate software and results.

https://doi.org/10.5281/zenodo.18111881

## Acknowledgments

I am grateful to Michael Lynch for the time he spent reviewing an early version of this manuscript and providing valuable feedback.

